# Growth parameters of the clupeids *Limnothrissa miodon* and *Stolothrissa tanganicae* in northern Lake Tanganyika (Bujumbura sub-basin)

**DOI:** 10.1101/2024.03.16.584994

**Authors:** T. N’sibula Mulimbwa, Cesc Gordó-Vilaseca, Leona J. M. Milec, Deepti M. Patel, Kankonda Busanga Alidor, Jean-Claude Micha, Jouko Sarvala, Joost A. M. Raeymaekers

## Abstract

The analysis of growth parameters in fish stocks, such as the asymptotic length (L_∞_) and the curvature parameter (K), is crucial for the estimation of production rates and total mortality rates in fisheries management. Here, we estimated the growth parameters of the clupeids *Limnothrissa miodon* and *Stolothrissa tanganicae* in the northern part of Lake Tanganyika (Bujumbura sub-basin). Both species are important targets of pelagic fisheries, but the available estimates are based on scarce size frequency data, and rarely on size as a function of age data. We provide new estimates of L_∞_ and K based on size as a function of age data, with age being inferred from otolith weights. Furthermore, we reanalyze several existing length frequency datasets using advanced statistical procedures to test if growth parameters estimated by length frequency analysis are consistent with those obtained from size as a function of age. We found that length frequency analysis consistently underestimates L_∞_ and overestimates K as compared to length-at-age data. We recommend that length-at-age data are collected at a much broader spatial and temporal scale than currently available to advance our understanding of the growth dynamics of these two sardine species. This will improve the assessment of stock productivity of Lake Tanganyika’s pelagic fisheries and the conservation of these economically important fish species.

## INTRODUCTION

Lake Tanganyika is about 10 million years old (Cohen *et al*. 1993) and is the second deepest and the fifth largest freshwater body in the world (Langenberg, 2008). So far, 1200 animal and plant species have been identified in this lake, ranking it among the most diverse lake ecosystems in the world. Among the main groups in this lake, fishes show a high degree of biodiversity (Cohen *et al*. 1993; Van Steenberge *et al*. 2011). Lake Tanganyika’s pelagic fisheries is a major source of food and economic activity of the human population living around the lake. These fish feed over 10 million people in the areas adjacent to Lake Tanganyika (Mölsä et al. 1999). Two small-sized clupeids, *Stolothrissa tanganicae* and *Limnothrissa miodon*, commonly known as “freshwater sardines”, along with the latid perch *Lates stappersii*, account for 60 % to 90 % of the total commercial catch in the pelagic environment (Mölsä et *al.* 1999). In the mid-1990s, annual harvests from Lake Tanganyika were estimated to range between 165 000-200 000 tons, quantities that translate into annual revenues of tens of millions of US dollars (Mölsä et *al.* 1999). More recent estimates are missing, because data coverage is spotty and there are long time gaps between monitoring projects (Van der Knaap 2014; Plisnier et al. 2023).

As for other African Great Lakes, the lack of a long-term monitoring programme for Lake Tanganyika creates uncertainty about the status of the lake and its pelagic fish stocks (Plisnier et al. 2023). Fisheries statistics published from the North of Lake Tanganyika suggest that annual catch rates (measured as catch per unit effort; CPUE) have been declining since the early 1980s (Mulimbwa 2006; Mulimbwa 2020; LTA Secretariat, 2012; Mushagalusa et al. 2015; Okito et al. 2017; Van der Knaap et al. 2014). The declines have been attributed to fishing pressure (Mulimbwa 2006; Mulimbwa 2020; LTA Secretariat, 2012; Mushagalusa et al. 2015; Okito et al. 2017; Van der Knaap et al. 2014), as well as to global warming (Plisnier, 1997; O’Reilly et al. 2003; Verburg et al. 2003). However, overfishing can only be assessed based on declines in the total annual catch, for which no reliable data exist (Van der Knaap et al. 2014). Interestingly, while fisheries stakeholders in the North of the Lake do indicate a noticeable decrease in fish abundance and size, they do not attribute these fishery problems to overfishing (De Keyzer et al. 2020). Yet another recent study found that CPUE has been stable in the North of Lake Tanganyika since 1993 (Sarvala & Mulimbwa 2023), indicating that not all fisheries statistics show the same trend. The impact of the effects of global warming on the pelagic fisheries has been questioned as well (Sarvala et al. 2006a, Sarvala et al. 2006b; Sarvala & Mulimbwa 2023). These uncertainties confirm that better fisheries monitoring is required to more accurately forecast fishing yields, assess fisheries potential, and understand the impact of fishing activities (Kolding et al. 2019).

One cornerstone of adequate fisheries management concerns the analysis of growth. Growth parameters such as the asymptotic length (L_∞_) and the curvature parameter (K) form the basis to estimate production rates and total mortality rates, which enable the prediction of fishing yields (Beverton & Holt 1957). Growth curves are used where age data are not available, as in most tropical fish stock assessments (Sparre & Venema 1989). L_∞_ and K can either be determined based on changes in length frequency distributions over time (length-frequency analysis; LFA) or based on length as a function of age data (length-at-age analysis; LAA). LFA has the advantage that size data are easy to obtain. However, the method requires young fish and frequent catch data to identify subsequent cohorts and track their modal growth progression. In environments and species with multiple periods of reproduction throughout the year, multiple overlapping cohorts and multimodal size frequency distributions may complicate cohort identification. Here, LAA might be more powerful. A commonly used technique for age determination in fish involves counting natural growth rings on the otoliths (Beamish & McFarlane 1983). In several tropical fishes, the diurnal transition from light to dark leaves a mark on the otoliths, and these daily increments can be used to estimate fish age in days (Kaningini, 1995; Kumura, 1995; Meisfjord et al. 2006; Chifamba 2019). This method is suitable for tropical environments where the temperature is almost constant throughout the year. While counting daily otolith rings can be time consuming, otolith radius and otolith weight tend to correlate well with the number of daily increments and are therefore also suitable to establish length-age relationships (Lou et al. 2005).

In this study, we estimated growth parameters for *S. tanganicae* and *L. miodon*. Various studies have gained useful insights in the biology of both clupeids, including their evolutionary origin (Milec et al. 2022), population genetic structure (De Keyzer et al. 2019; Junker et al. 2020), position in the food web (Sarvala et al. 1999; Ehrenfels et al. 2023), parasites (Kmentová et al. 2020), reproductive biology (Mulimbwa et al. 2014a; Mulimbwa et al. 2022), and responses to limnological changes (Kimirei & Mgaya, 2007; Plisnier et al. 2009; Mulimbwa et al. 2014b). While *S. tanganicae* is fully pelagic, *L. miodon* makes more use of the littoral areas, also for reproduction and as nursery grounds (Mulimbwa et al. 2022). Both fish species are the target of artisanal fisheries in the pelagic, but in shallow littoral waters, juveniles of *L. miodon* are also subjected to the relatively low selectivity of beach seines (Mölsä *et al*. 2002). Basic information for fisheries management, such as age at maturity and size at maturity are available for both species, although there are no in-depth studies assessing the variability in those parameters. In addition, the data are patchy in both time and space. Age at maturity in Lake Tanganyika is usually estimated at 8.5 months corresponding to a total length of 78 mm for females and 77 mm for males in *S. tanganicae*, and at 8.6 months corresponding to a total length of 101 mm for females and 95 mm for males in *L. miodon* (Mannini 1998). Growth parameters have been obtained for both species as well (Chapman & van Well, 1978; Roest, 1978; Coulter, 1991; Moreau et al. 1991; Mulimbwa & Shirakihara, 1994; Kimura 1995; Ahonen 2001; Mulimbwa 2006). Most studies have inferred growth parameters using LFA on length–frequency distributions from catch data. The few exceptions using LAA include Kimura (1991b, C), Kimura (1995) and Ahonen (2001). Especially in *S. tanganicae*, the most abundant cohort is often the one recruited during the dry season (Mulimbwa et al. 2014b). Yet, multiple partially overlapping cohorts do occur in both species, and pose challenges of cohort delineation for the assessment of growth.

To address the need for more accurate growth data and validate existing growth data for the two clupeid fish species, we provide new estimates of L_∞_ and K based on length-at-age data, with age inferred from otolith weight. Furthermore, we reanalyze three published datasets with length frequency data using more recent and advanced statistical procedures, and test if growth parameters estimated by size frequency analysis are consistent with those obtained from length-at-age data.

## MATERIAL AND METHODS

### Study design

This study combines datasets from different surveys on the two clupeids that were all conducted in the northern end of Lake Tanganyika (03° 28’ S, 29° 17’ E; Fig. 1) in the Bujumbura sub-basin. We included (1) length frequency data for the period 1987-1989 from Mulimbwa and Shirakihara (1994), (2) length frequency data for the period 1999-2001 from Mulimbwa (2006), (3) length as a function of age data for the period 2004-2005 from Mulimbwa et al. (2014b), and length frequency data for the period 2007-2008 from Mulimbwa et al. (2014b) (Table 1). Sampling and data collection procedures for each data type are described in more detail below.

**Figure 1.**
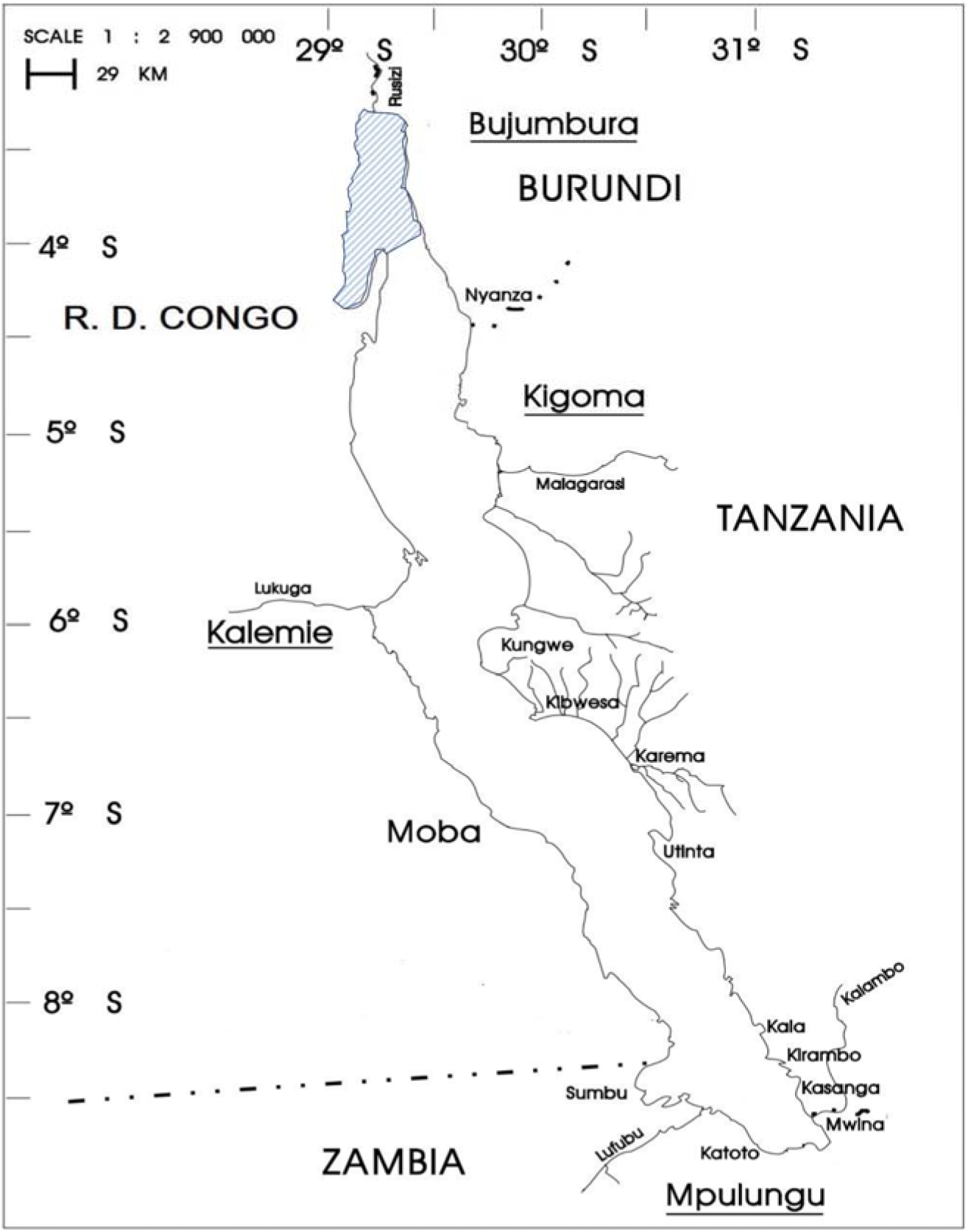
Geographic position of Lake Tanganyika bordered by DR Congo, Burundi, Tanzania and Zambia. The shaded area marks the Bujumbura sub-basin.

**Table 1.** Overview of datasets used in this study for the analysis of growth parameters including survey period, total sample size (N), data type (LF: length frequency data; LA: length-at-age data), published estimates of L_∞_ (TL in cm) and K, and reference. NA indicates that no information is available from the original publications. Estimates of L_∞_ marked with an asterisk are converted from SL to TL following Mannini *et al*. (1996).

| Survey period (months) | N | Data type | $L_{\infty}$ | K | Reference |
| --- | --- | --- | --- | --- | --- |
| *S. tanganyicae* |  |  |  |  |  |
| Nov 1987 - Apr 1989 (18) | 24,167 | LF | 12.34* | 2.14 | Mulimbwa & Shirakihara 1994 |
| May 1999 - Nov 2001 (30) | 84,503 | LF | 11.70 | 2.6 | Mulimbwa 2006 |
| 2004 - 2005 | 260 | LA | NA | NA | Mulimbwa et al. 2014b |
| Feb 2007 - May 2008 (16) | NA | LF | NA | NA | Mulimbwa et al. 2014b |
| *L. miodon* |  |  |  |  |  |
| Nov 1987 - Apr 1989 (18) | 3,834 | LF | 16.12* | 1.15 | Mulimbwa & Shirakihara 1994 |
| May 1999 - Nov 2001 (27) | 12,272 | LF | 17.90 | 0.74 | Mulimbwa 2006 |
| 2004 - 2005 | 244 | LA | NA | NA | Mulimbwa et al. 2014b |
| Feb 2007 - May 2008 (16) | NA | LF | NA | NA | Mulimbwa et al. 2014b |

As *S. tanganicae* is entirely pelagic, all data for this species originate from samples collected in the pelagic environment. In contrast, *L. miodon* occurs in the pelagic environment as well as in littoral areas of the lake. The data for *L. miodon* were obtained from both the littoral and the pelagic in 1987 – 1988 and in 1999 – 2001, while in 2004 – 2005 and in 2007-2008, the species was only sampled in the pelagic.

Because the available length frequency data (1987-1989, 1999-2001, 2007-2008) and length at age data (2004-2005) originated from different survey periods, our study design is not optimal to compare growth parameters based on the two data types. Nevertheless, since the length-at-age survey period is embedded within the two decades spanning the three length frequency surveys, it should still be possible to distinguish effects of data type from temporal patterns, especially when temporal variation is minor.

### Length-at-age data

Briefly, 260 specimens of *S. tanganicae* and 244 specimens of *L. miodon* were collected in the pelagic from the study area in 2004-2005 (Table 1). First, the total length (TL) of each specimen was measured to the nearest mm, and otoliths were extracted and dried at room temperature at the CRH laboratory in Uvira (DR Congo). Subsequently, the otoliths were weighed with a microbalance (µg; dry weight at 20 °C) at the University of Turku (Finland). Finally, the age of these fish was estimated using the relationship between otolith dry weight (ODW) in µg and age in days based on counts of daily rings following the formula age = (ODW + a)/b, where a = 99.4, b = 2.42 for *S. tanganicae* and a = 120.8, b = 2.53 for *L. miodon* (Ahonen 2001). Ahonen (2001) indicated that the relationship between ODW and age was linear or near-linear, with a high Pearson correlation coefficient in both species (*S. tanganicae*: R^2^ = 0.88; *L. miodon*: R^2^ = 0.96). Simple linear regressions of ODW versus age for the two species, as originally established by Ahonen (2001), are visualised in Supplementary Figure 1.

### Length frequency data

For all three studies, samples were taken from artisanal fishing units using lift nets with a mesh size of 4.4 mm (knot to knot) in the upper four-fifths of the net and 3.3 mm (knot to knot) for the rest of the net. The samples were collected twice a week during the investigation period. After collection, samples were sorted by species at the CRH laboratory in Uvira (DR Congo). Their total lengths (TL) were determined to the nearest mm, and the specimens were grouped into size classes of 2 mm for *S. tanganicae*, and 4 mm for *L. miodon*, in 1999-2001 and 2007-2008. The data from 1987-1989 were collected in size classes of 5 mm for both species. Total sample sizes for each period are shown in Table 1.

For 1987 – 1988, the original data from Mulimbwa and Shirakihara (1994) could not be retrieved from the authors. Instead, we digitized the length frequency plots in Figure 2 (*S. tanganicae*) and Figure 4 (*L. miodon*) in Mulimbwa and Shirakihara (1994) using WebPlotDigitizer version 4.6 (Rohatgi 2022). Since length in the 1987-1988 study was measured in mm as standard length (SL), size classes were converted to total length (TL) in mm following the formula TL = a + (b * SL), where a = 0.8729, b = 1.1562 for *S. tanganicae* and a = 1.6658, b = 1.1873 for *L. miodon* (Mannini *et al*. 1996).

**Figure 2.**
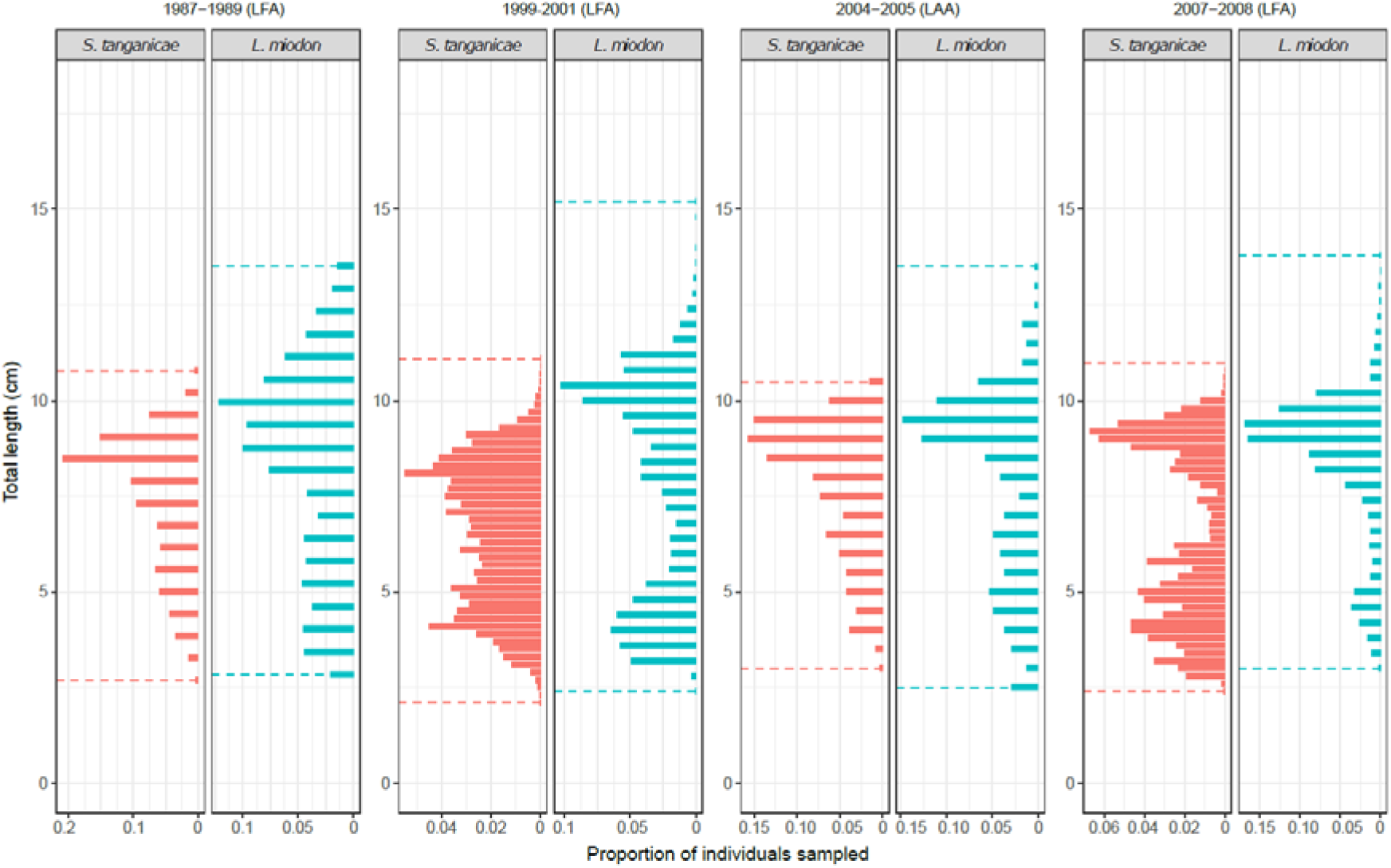
Total length distribution (cm) for both species and the four survey periods.

### Data analysis

#### 1) Length-at-age analysis

A Von Bertalanffy (1938) growth function (VBGF) was fitted to the otolith derived length-age data, and its parameters L_∞_ (cm) and K (year^-1^) were determined using the non-linear least squares method, available within the function “growth_length_age()” in the TropFishR package (Mildenberg et al. 2017).

#### 2) Length frequency analysis

The growth parameters to fit a VBGF to length frequency data have historically been estimated using several methods (e.g. the Power-Wetherall or Ford-Walford method), most of which have been questioned for containing intrinsic biases from different sources (Schwamborn 2018). However, recent improvements to ELEFAN (electronic length frequency analysis, Pauly & David 1981), the most widely used method for fitting a growth curve to length frequency data, provide a promising new path for length-based studies of aquatic populations. These advancements include the estimation of both L_∞_ and K with fewer assumptions than previous models (Taylor & Mildenberg 2017), as well as the calculation of their associated uncertainties using bootstrapping (Schwamborn et al. 2019). We applied these new methods using the TropFishR and fishboot packages in R (Mildenberg et al. 2017; Taylor & Mildenberg 2017; Schwamborn et al. 2019) with the parameters shown in Table 2.

**Table 2.** Parameters used in the VBGF fitting using ELEFAN bootstrapping in each of the three surveys based on length frequency data. GA: Genetic algorithm; SA: Simulated annealing; MA: moving average; L_∞_: asymptotic total length; K: annual growth coefficient; t_anchor: peak spawning month; nresamp: number of permutations.

| Species | Algorithm | Parameter (unit) | 1987-1988 | 1999-2001 | 2007-2008 |
| --- | --- | --- | --- | --- | --- |
| *S. tanganyicae* | GA, SA | MA (bins) | 5 | 7 | 5 |
| | | $L_{\infty}$ (cm) | 6-12 | 6-12 | 6-12 |
|  |  | K (year<sup>-1</sup>) | 1-10 | 1-10 | 1-10 |
|  |  | t_anchor (year) | 0-1 | 0-1 | 0-1 |
|  |  | nresamp | 400 | 400 | 400 |
|  |  | Agemax (year) | 18/12 | 18/12 | 18/12 |
|  |  | bin_size (cm) | 0.8 | 0.8 | 0.8 |
| *L. miodon* | GA, SA | MA (bins) | 5 | 7 | 7 |
| | | $L_{\infty}$ (cm) | 8-22 | 8-22 | 8-22 |
|  |  | K (year<sup>-1</sup>) | 1-10 | 1-10 | 1-10 |
|  |  | t_anchor (year) | 0-1 | 0-1 | 0-1 |
|  |  | nresamp | 400 | 400 | 400 |
|  |  | Agemax (year) | 30/12 | 30/12 | 30/12 |
|  |  | bin_size (cm) | 0.8 | 0.8 | 0.8 |

ELEFAN relies on attributing different scores to restructured different length bins, depending on how much each bin deviates from a moving average across a user-defined number of length bins. This was done by grouping the size classes into bins of 0.8 cm each, and a moving average of 5-7 size classes (Table 2), depending on the size of the smallest cohort, following the general rule of thumb (Taylor & Mildenberg 2017). Afterwards, a score is given to different growth curves, depending on whether they intersect with positively or negatively scored length bins (Gayanilo & Pauly 1997). The curve with the highest score is considered the best fitting curve to the data. Finding the best fitting curve can be computationally costly. Two optimization approaches (“simulated annealing” and “genetic algorithm”) have been implemented for growth function fitting in R (Taylor & Mildenberg 2017; Mildenberg et al. 2017). Both optimization approaches can find high-scoring solutions, but because several models often score high, different solutions can be reached depending on the initial settings, which can create an intrinsic source of bias. To overcome this limitation, bootstrapping methodology can be applied, fitting a large number of growth curves with similar scoring solutions, and finally calculating the uncertainty of the mean value of the estimations (Schwamborn et al. 2019). We estimated L_∞_ and K of both species from length frequency data by bootstrapping both algorithms, using the package “fishboot” in R (Schwamborn et al. 2019), and we report the probability distribution of the estimations. The total parameter space considered ranged from 1 to 10 for K for both species (Table 2). For L_∞_, we considered values from 6 cm to 12 cm for *S. tanganicae*, and from 8 to 22 cm for *L. miodon* (Table 2). Here, the upper values are above the maximum total length reported for each species in Fishbase (*S. tanganicae*: 10 cm SL ∼ 11.65 cm TL; *L. miodon*: 17 cm SL ∼ 20.35 cm TL; Froese & Pauly 2017). We manually set a maximum age to each species (Table 2; 18 months for *S. tanganicae* and 30 months for *L. miodon*; Mannini et al. 1996), which is a key step for the calculation of the growth parameters in this method.

#### 3) Consistency between methods

To assess the consistency between length-at-age-based and size frequency-based methods, the agreement of growth parameters (L_∞_ and K) between different sampling periods was tested by comparing the 95 % confidence intervals of the estimates based on LAA with the 95 % credibility intervals of the estimates based on LFA.

## RESULTS

### S. tanganicae

Length frequency data across time periods ranged between 2.1 and 11.1 cm TL. Recruitment was continuous throughout the year, as indicated by small individuals (2.1 - 4.0 cm) appearing all year round in the length frequency data of the three survey periods. Length-at-age data comprised individuals ranging between 3.3 cm and 10.6 cm in TL, and between 56 and 372 days in age. The length distribution of the length frequency data and the length-at-age data was comparable (Fig. 2).

Based on length-frequency data analysis (LFA), estimates of L_∞_ (range: 8.38 - 9.56 cm) were consistent between estimation methods with narrow confidence intervals, and showed substantially lower values in 1987-1989 and 1999-2001 than in 2007-2008 (Table 3; Supplementary Figure 2). In contrast, the annual K (range: 5.48 – 8.46) showed high uncertainty which led to an overlap in confidence intervals between most study periods and estimation methods (Table 3). Based on the length-at-age data (LAA) from 2004-2005, we obtained an estimate of 10.64 cm for L_∞_ and 3.18 for K (Table 3). Non-overlapping credibility and confidence intervals indicated that all estimates of L_∞_ and most estimates of K were respectively significantly lower and higher for LFA than for LAA. According to the LAA curve, *S. tanganicae* grows rapidly during the first five months of its life and growth becomes almost constant after ten months (Fig. 3; Fig. 4; Supplementary Figure 3).

**Figure 3.**
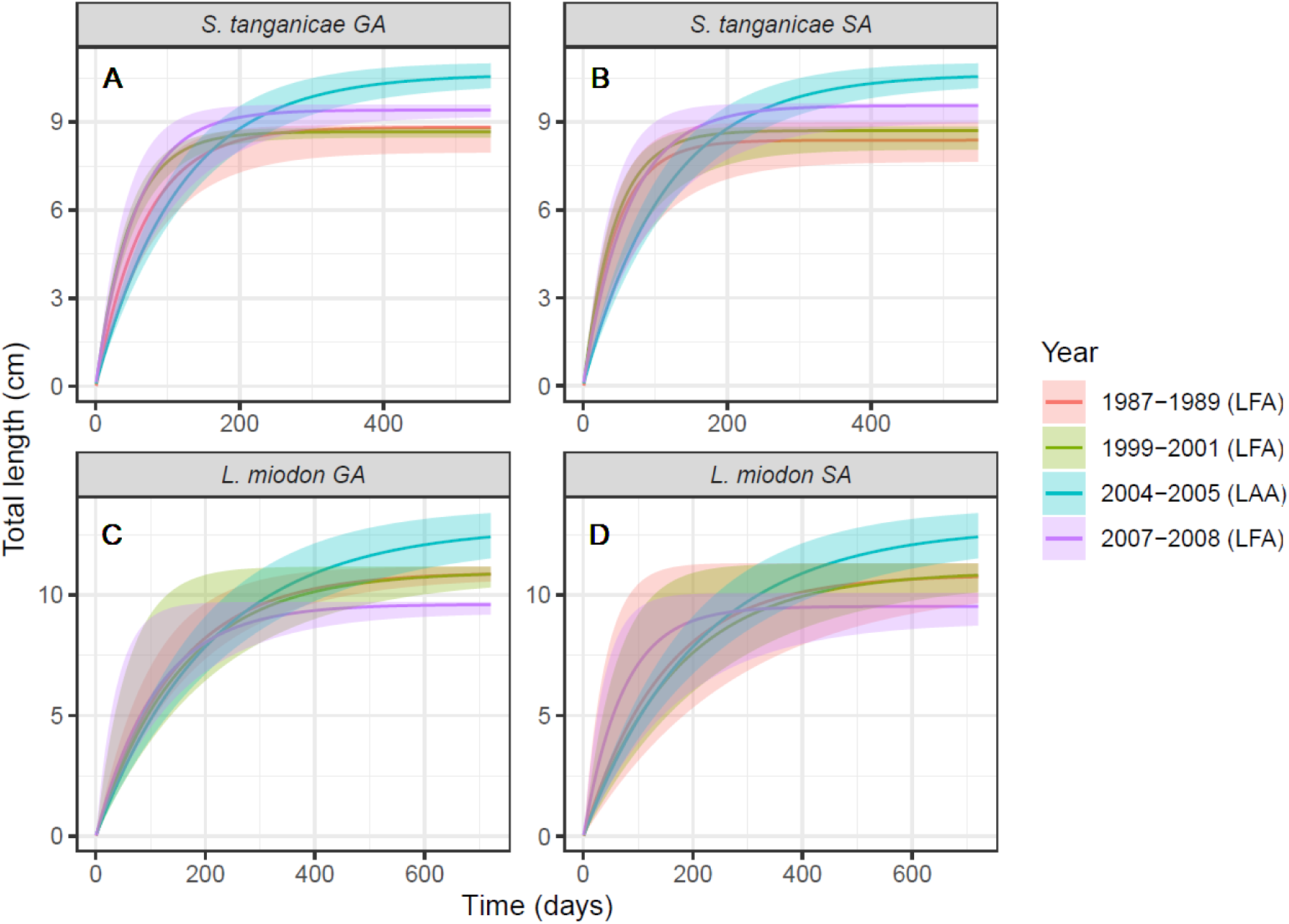
VBGF estimated from total length frequency data of 1987-1988, 1999-2001 and 2007-2008 for (A) *Stolothrissa tanganicae* with Genetic Algorithm (GA); (B) *Stolothrissa tanganicae* with Simulated Annealing (SA); (C) *Limnothrissa miodon* with GA; and (D) *Limnothrissa miodon* with SA. Each plot is superimposed with VBGF estimates from otolith-derived length at age data of 2004-2005, along with its 95 % confidence interval. Bayesian credibility intervals (in the case of LFA) or confidence intervals (in the case of LAA) were obtained from growth curve estimations for the upper and lower limits of the 95 % credibility or confidence intervals of parameters L_∞_ and K.

**Figure 4.**
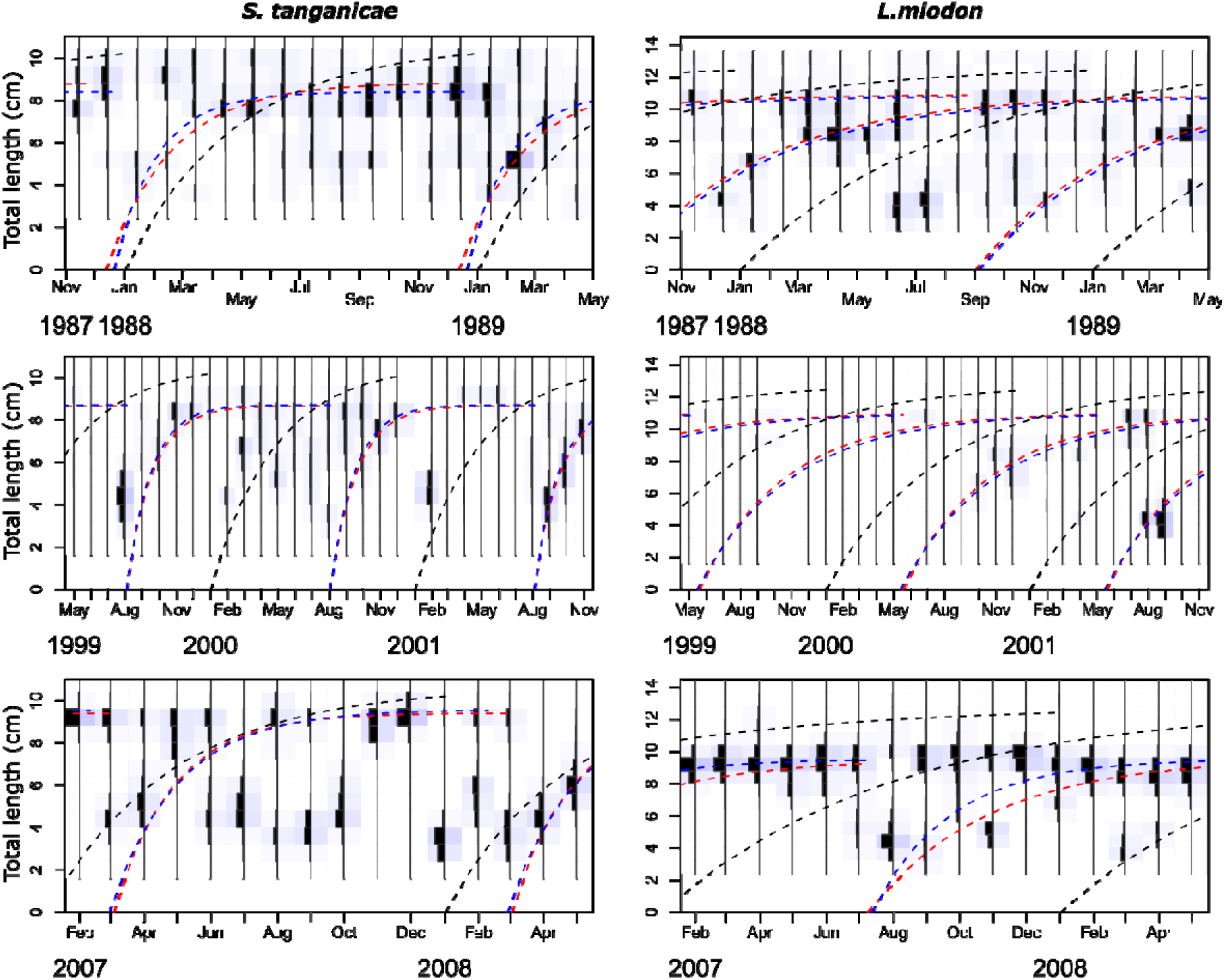
Total length frequency data and corresponding Von Bertalanffy growth curves for the three survey periods for *Stolothrissa tanganicae* (left panels) and *Limnothrissa miodon* (right panels). The red and blue dashed lines mark growth curves estimated with the genetic algorithm and the simulated annealing algorithm, respectively. For comparison, the growth curves estimated from length-at-age data are superimposed (black dashed lines with t_0_ set to 1^st^ of January). The length frequency data are plotted in dark blue, and blue shades reflect the frequency of each length class.

**Table 3.** Growth parameters of clupeids (*S. tanganicae*, *L. miodon*) estimated for different time periods and using different estimation procedures for length frequency analysis (LFA) and length-at-age analysis (LAA). Estimation methods include two optimization approaches in the case of LFA (SA: ELEFAN with simulated annealing; GA: ELEFAN using genetic algorithm), and the non-linear least-squares method (LSM) in the case of LAA. L_∞_: asymptotic total length (cm), K: annual growth coefficient. CI: Bayesian credibility intervals (in the case of LFA) or confidence intervals (in the case of LAA).

| Survey period | Analysis | Estimation | $L_{\infty}$ [95 % CI] | K [95 % CI] |
| --- | --- | --- | --- | --- |
| *S. tanganyicae* |  |  |  |  |
| 1987-1989 | LFA | GA | 8.82 [7.97-8.90] | 5.48 [4.49-7.59] |
|  |  | SA | 8.38 [7.65-8.99] | 8.27 [4.66-9.91] |
| 1999-2001 | LFA | GA | 8.67 [8.47-8.80] | 7.86 [6.81-8.79] |
|  |  | SA | 8.71 [8.06-8.84] | 8.46 [5.01-10.42] |
| 2004-2005 | LAA | LSM | 10.64 [10.32-11.05] | 3.18 [2.76-3.63] |
| 2007-2008 | LFA | GA | 9.41 [9.20-9.60] | 6.64 [3.72-9.35] |
|  |  | SA | 9.56 [9.01-9.64] | 5.89 [3.57-9.99] |
| *L. miodon* |  |  |  |  |
| 1987-1989 | LFA | GA | 10.93 [10.72-11.18] | 2.55 [2.12-3.63] |
|  |  | SA | 10.83 [10.46-11.31] | 2.49 [1.30-9.42] |
| 1999-2001 | LFA | GA | 10.96 [10.69-11.17] | 2.36 [1.69-6.34] |
|  |  | SA | 10.97 [10.65-11.30] | 2.16 [1.51-6.13] |
| 2004-2005 | LAA | LSM | 12.82 [12.18-13.64] | 1.73 [1.47-2.01] |
| 2007-2008 | LFA | GA | 9.61 [9.26-9.70] | 3.30 [2.44-9.84] |
|  |  | SA | 9.52 [8.87-10.07] | 5.07 [2.10-10.09] |

### L. miodon

Length categories of the length-frequency data across time periods ranged between 2.4 and 15.2 cm TL. Recruitment occurred less frequently throughout the year than in *S. tanganicae*, as small cohorts (2.4 - 6.0 cm) were less frequent across the three survey periods. Length-at-age data comprised individuals ranging between 2.5 cm and 13.6 cm in TL, and between 62 and 571 days in age. The length distribution of the length frequency data and the length-at-age data was comparable (Fig. 2).

Based on length-frequency data, estimates of L_∞_ (range: 9.52 - 10.97 cm) were consistent across estimation methods, but were lower in 2007-2008 than in 1987-1989 and 1999-2001 (Table 3; Supplementary Figure 2). Ranging between 2.16 and 5.07, the annual K was more consistent across the three LFA survey periods and estimation methods than estimations for *S. tanganicae*, although the uncertainty was generally also high (Table 3). Based on the length-at-age data from 2004-2005, we obtained an estimate of 12.82 cm for L_∞_ and 1.73 for K (Table 3). Similar to *S. tanganicae*, non-overlapping credibility and confidence intervals intervals indicated that all estimates of L_∞_ and half of the estimates of K were respectively significantly lower and higher for LFA than for LAA. According to the LAA curve, *L. miodon* grows rapidly during the first ten months of its life and growth becomes almost constant after 18 months (Fig. 3; Fig. 4; Supplementary Figure 3).

## DISCUSSION

The objective of this study was to examine the growth patterns of *S. tanganicae* and *L. miodon* in the North part of Lake Tanganyika. We compared growth estimates derived from observed length frequency data and indirectly estimated length-at-age data, collected over different survey periods. This comparison informs us about the accuracy of size frequency-based estimation methods for growth parameters, and the necessity of complementing fisheries management with more complete life history information using age-based data. For both species, we consistently found lower estimates of L_∞_ and higher estimates of K for length frequency data than for length at age data. Below we discuss the reliability of this result, and its implications for the management of the artisanal fisheries.

Because length frequency and length at age data were collected in different survey periods, estimation procedures and survey periods are confounded in our study, and therefore do not allow us to directly compare the results. Nevertheless, because the differences in estimates between the two procedures largely exceed the differences between survey periods within LFA, it is unlikely that temporal variation alone can account for the observed differences between the two methods. It is indeed more likely that the disparities between data types and estimation methods stem from methodological nuances that influence the estimation of growth parameters. Otolith-based methods are believed to provide accurate estimations of age, which together with direct measurements of length, should provide accurate growth curves (Schwamborn et al. 2023). Ahonen (2001) demonstrated that the age of the two sardine species can be reliably estimated from otolith dry weight (Pearson correlation between otolith dry weight and age based on counts of daily otolith rings: r^2^ = 0.88 for *S. tanganicae*; r^2^ = 0.96 for *L. miodon*; Supplementary Figure 1). While this relationship was established using individuals from the central part of Lake Tanganyika (Kigoma, Malagarasi and Kipili; Ahonen 2001), we applied it here to individuals from the North of Lake Tanganyika. Nevertheless, this should yield acceptable results, and overall, the accuracy of our LAA-based estimates most likely exceeds the accuracy of our LFA-based estimates.

In general, LFA is highly sensitive to sampling bias, including the lack of large, old individuals (Schwamborn 2018; Schwamborn et al. 2019; Wang et al. 2021), as well as the absence of young cohorts. Other potential sources of sampling bias include the concentration of fishing efforts on commercially valuable areas, and differences in fishing gears and methods. These potential issues with LFA likely apply to the two Lake Tanganyika sardine species, as notified previously by Mulimbwa & Shirakihara (1994). In the present study, the discrepancy between LFA-based and LAA-based estimates may be due to an underrepresentation of the largest individuals in the length-frequency data of both species. The relatively low number of young cohorts in *L. miodon* may also have lowered the precision of LFA in this species. Avoiding sampling bias for size is particularly challenging in *L. miodon* because large, mature individuals are more commonly observed in littoral areas than in the pelagic zone, so the sampling effort in each area can directly affect the results (Mulimbwa & Shirakihara 1994; Mulimbwa et al. 2022). Since no *L. miodon* were collected in the littoral areas during the most recent survey period (2007-2008), part of the variation in LFA-based estimates of L_∞_ and K between years might be due to sampling bias. However, similar fluctuations in LFA-based estimates of L_∞_ and K were observed in *S. tanganicae*, which was always sampled in the pelagic. Considerable variation in growth curves based on LFA is thus expected. Currently, it remains unclear if LAA data generate more consistent results than LFA. Our LAA-based estimates of L_∞_ for both *S. tanganicae* and *L. miodon* from catches in 2004-2005 were substantially lower than the LAA-based estimate for both species from catches in 1990 by Kimura (1995) (Figure 5). However, both studies were conducted at opposite ends of the lake, and spatial variation in environmental conditions may also affect growth trajectories. Overall, better data are required to assess the robustness of growth parameters.

**Figure 5.**
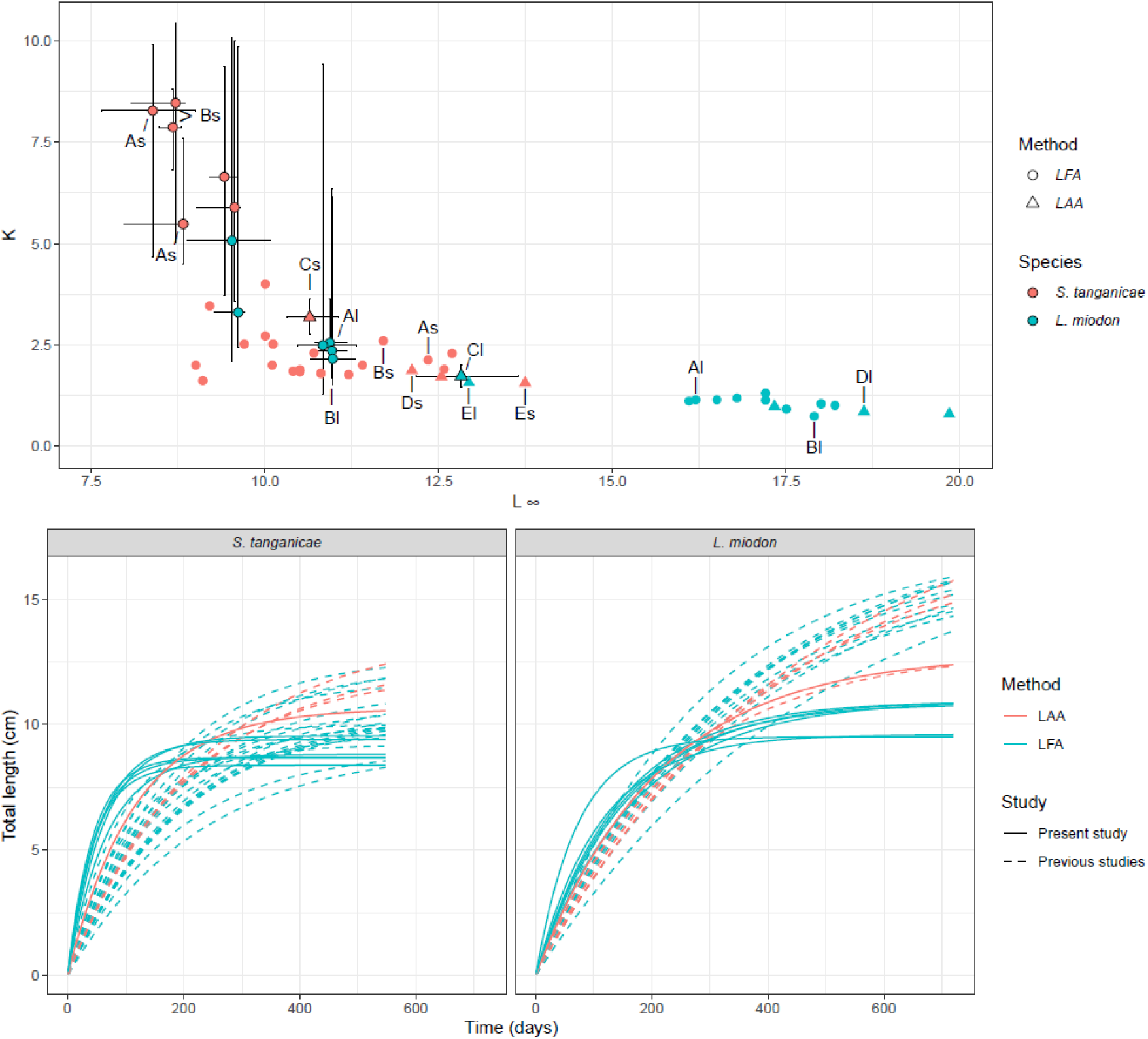
**Top**: annual growth coefficient (K) against asymptotic total length (L_∞_ in cm) for *S. tanganicae* (red) and *L. miodon* (blue) based on length-frequency analysis (LFA; circles) or length-at-age analysis (LAA; triangles). Data points with black edges and error bars (95 % confidence intervals) are generated by this study and are listed in Table 3. Data points without error bars are derived from the overview provided in Table 9 and Figure 47 in Mannini et al. (1996), in addition to estimates by Ahonen (2001) and Mulimbwa (2006). Labeled data points are based on LFA data by Mulimbwa & Shirakihara (1994) (As: *S. tanganicae*; Al: *L. miodon*), LFA data by Mulimbwa (2006) (Bs: *S. tanganicae*; Bl: *L. miodon*), LAA data by Mulimbwa et al. (2014b) (Cs: *S. tanganicae*; Cl: *L. miodon*), LAA data by Kimura (1995) (Ds: *S. tanganicae*; Dl: *L. miodon*), and LAA data by Ahonen (2001) (Es: *S. tanganicae*; El: *L. miodon*). **Bottom**: Von Bertalanffy growth curves for the present study and previous studies by species (*Stolothrissa tanganicae* vs. *Limnothrissa miodon*) and method (LAA vs. LFA). Each curve corresponds to a data point of the top panel.

Another challenge of LFA is the aggregation of individuals into length bins, which has the potential to obscure the age structure within each bin (Taylor & Mildenberger 2017; Schwamborn, 2018). This can manifest as well in lower L_∞_ values and higher K values, particularly with bigger bin sizes (Wang et al. 2020). However, a recent study directly comparing LFA and LAA in lane snapper (*Lutjanus synagris*) found no difference in L_∞_ and K between the two methods (Schwamborn et al. 2023). Interestingly, Kimura (1995) concluded that the growth parameters obtained from length-at-age data (based on daily counts of otolith rings) generally agreed with those previously estimated from analyses of length– frequency distributions. However, close inspection of Kimura’s (1995) results reveals that the author’s otolith-based estimates for L_∞_ almost always exceeded the estimates for L_∞_ from length-frequency studies, and *vice versa* for K, and thus seemed to be subjected to the same bias we report here.

Finally, both our LAA-based and LFA-based estimates for the growth trajectory of the two sardine species from the North of Lake Tanganyika deviated substantially from estimates of previous studies (Table 1; Figure 5). This includes LAA-based and LFA-based studies from various regions of Lake Tanganyika (Figure 5), but also the length frequency datasets from the North of Lake Tanganyika for which we re-estimated the growth parameters (Figure 5; Table 1; Mulimbwa & Shirakihara 1994; Mulimbwa 2006). Almost all studies reported significantly larger values of L_∞_, and therefore lower values of K than the present study (Figure 5). This was true for both species, although the discrepancy for K between previous studies and the present study was more pronounced in *S. tanganicae*, while the discrepancy for L_∞_ was more pronounced in *L. miodon* (Figure 5). For the LFA-based estimates, this discrepancy could be explained by the fact that previous studies are largely based on outdated methods such as the Ford-Walford plot, which are known to generate inaccurate results (Schwamborn 2018). Indeed, the improved ELEFAN methods we used here are based on fewer assumptions and allow for screening a large range of parameter values, which should lead to more precise and easily reproducible results (Mildenberger et al. 2017; Schwamborn et al. 2019). Still, the subjective choice of suitable parameter space for L_∞_ can often result in biased results with narrow confidence intervals (Schwamborn et al. 2023). For this reason, we opted for upper parameter limits above the maximum length reported in Fishbase for each species, which should result in unbiased estimations with realistic uncertainty estimates. As a result, some of our LFA-based estimates, especially for K, come with very high uncertainties, which may indicate that the quality of the available length frequency data is insufficient to achieve robust conclusions. This leaves little room for scientific advice beyond improving monitoring efforts to obtain better and more accurate growth parameters based on LFA.

### Conclusions and implications for fisheries management

Growth parameters link age and size and are thus key to all length-based stock assessment models where data on catches by age are not available. The lower L_∞_ and higher K values obtained with length frequency data relative to length-at-age data for the two sardine species emphasizes the need for future research aimed at advancing our understanding of growth dynamics and their implications for the pelagic fisheries of Lake Tanganyika. We recommend that length-at-age data are obtained at a much broader spatial and temporal scale than is currently available. However, as length frequency data can be more easily collected, such data will remain our primary source of information and should continue to be obtained. Furthermore, both length frequency data and length-at-age data should be collected for the same study periods to test for a systematic bias between the two methods. In addition, it is important that all samples in a time series come from the same landing site and fishing area to rule out that changes in length frequencies over time are due to sampling bias. As we have shown, it is straightforward to reanalyze existing length frequency data whenever the original data can be retrieved, but it is also possible to regenerate the data after digitizing published length frequency plots.

## ACKNOWLEDGEMENTS

We thank Jeppe Kolding and one anonymous reviewer for critical feedback. This study was partially funded by the Research Council of Norway through project 343414 “CHIPOLATA - Climate change and harvesting impacts on the pelagic fish community of Lake Tanganyika”.

## DATA AVAILABILITY

Datasets including the two files with length-at-age data and the six files with length frequency data, as well as the script to run the analysis with the TropFishR package in R are available on Zenodo (https://doi.org/10.5281/zenodo.12812204) and Github (https://github.com/CescGV/Parameters_Limno_Stolo_Tanganyika-).

## SUPPLEMENTARY FIGURES

**Supplementary Figure 1.**
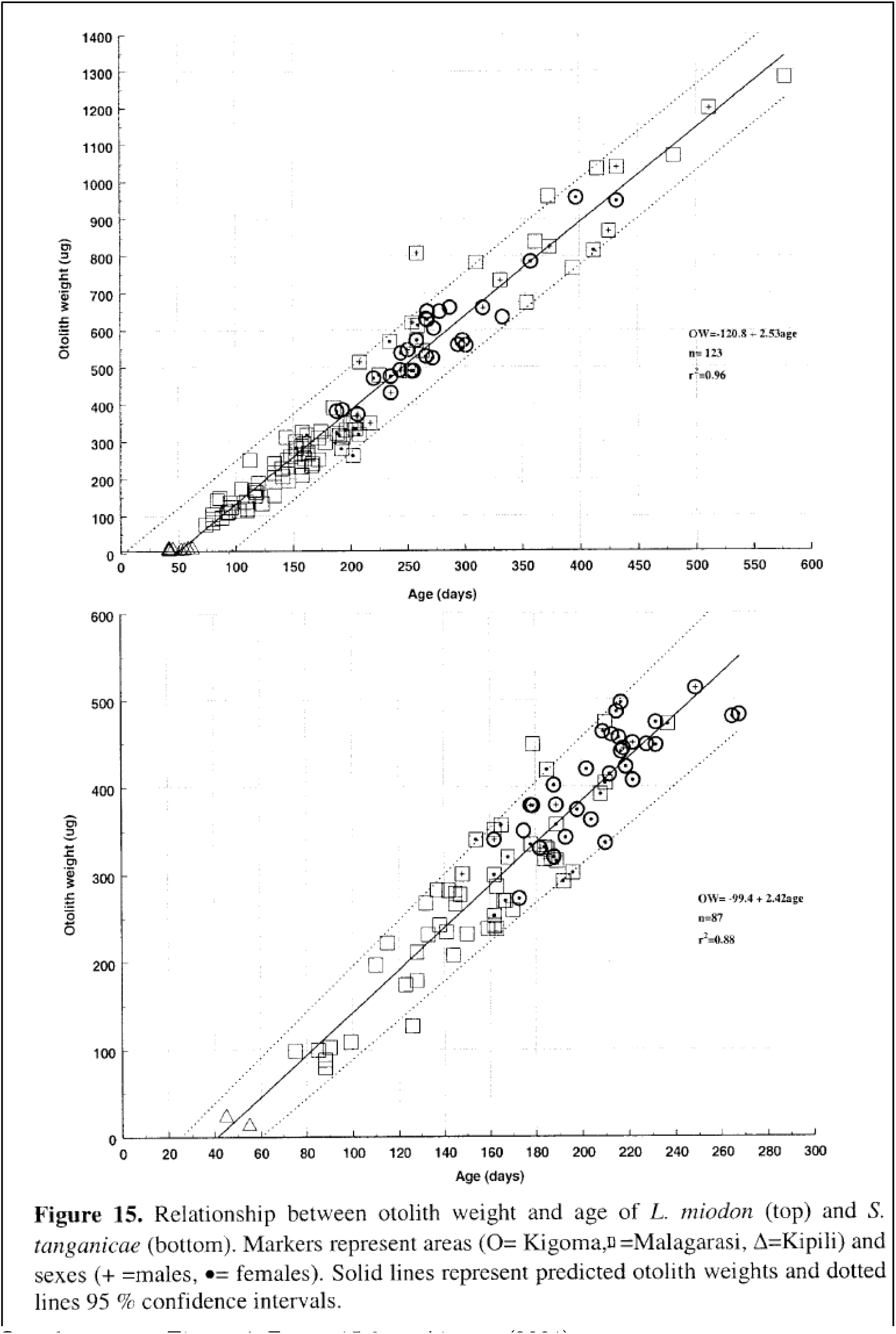
Figure 15 from Ahonen (2001).

**Supplementary Figure 2.**
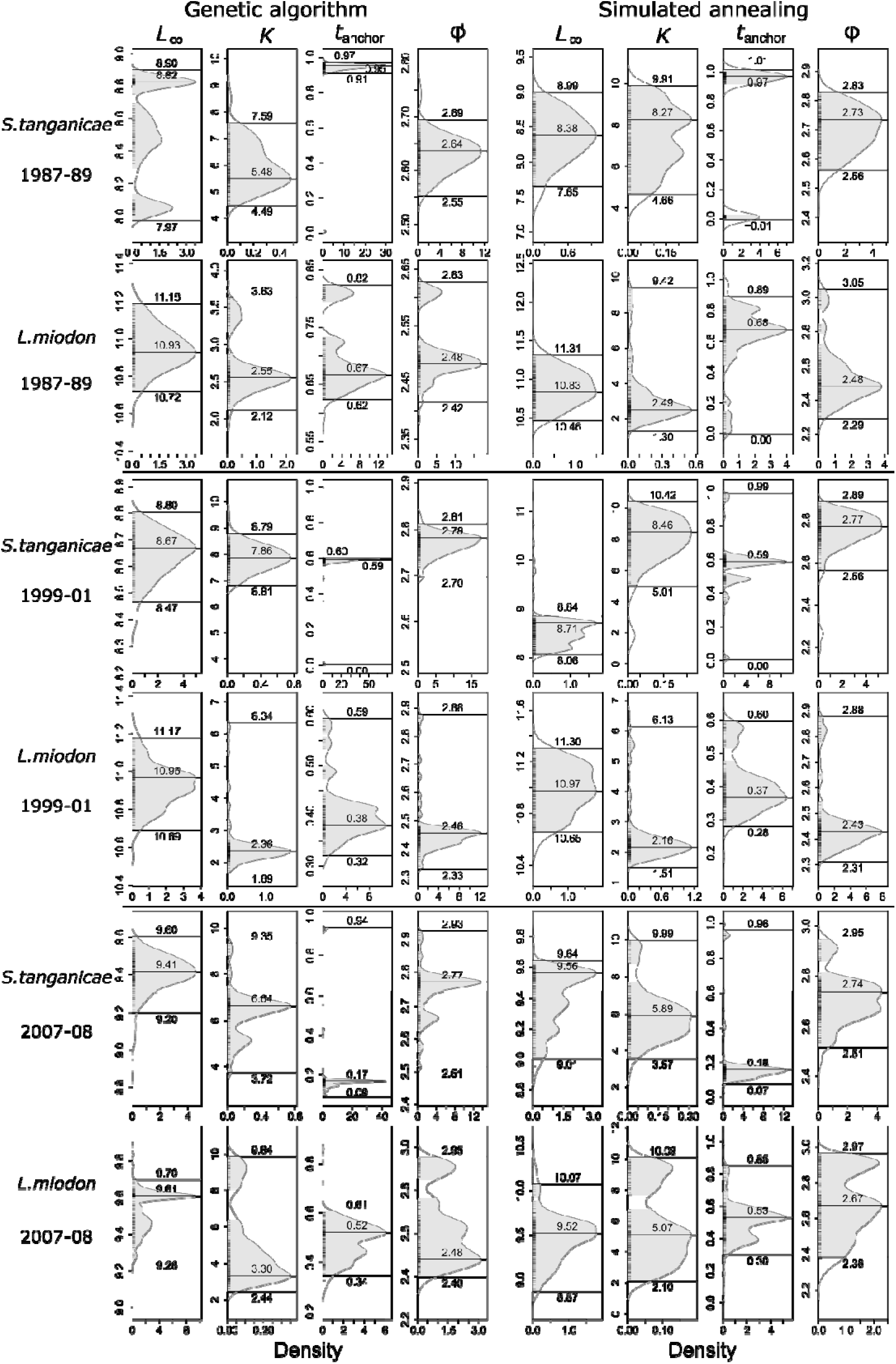
Probability distribution of estimated parameters using genetic and simulated annealing algorithms for *Stolothrissa tanganicae* and *Limnothrissa miodon* in each study period.

**Supplementary Figure 3.**
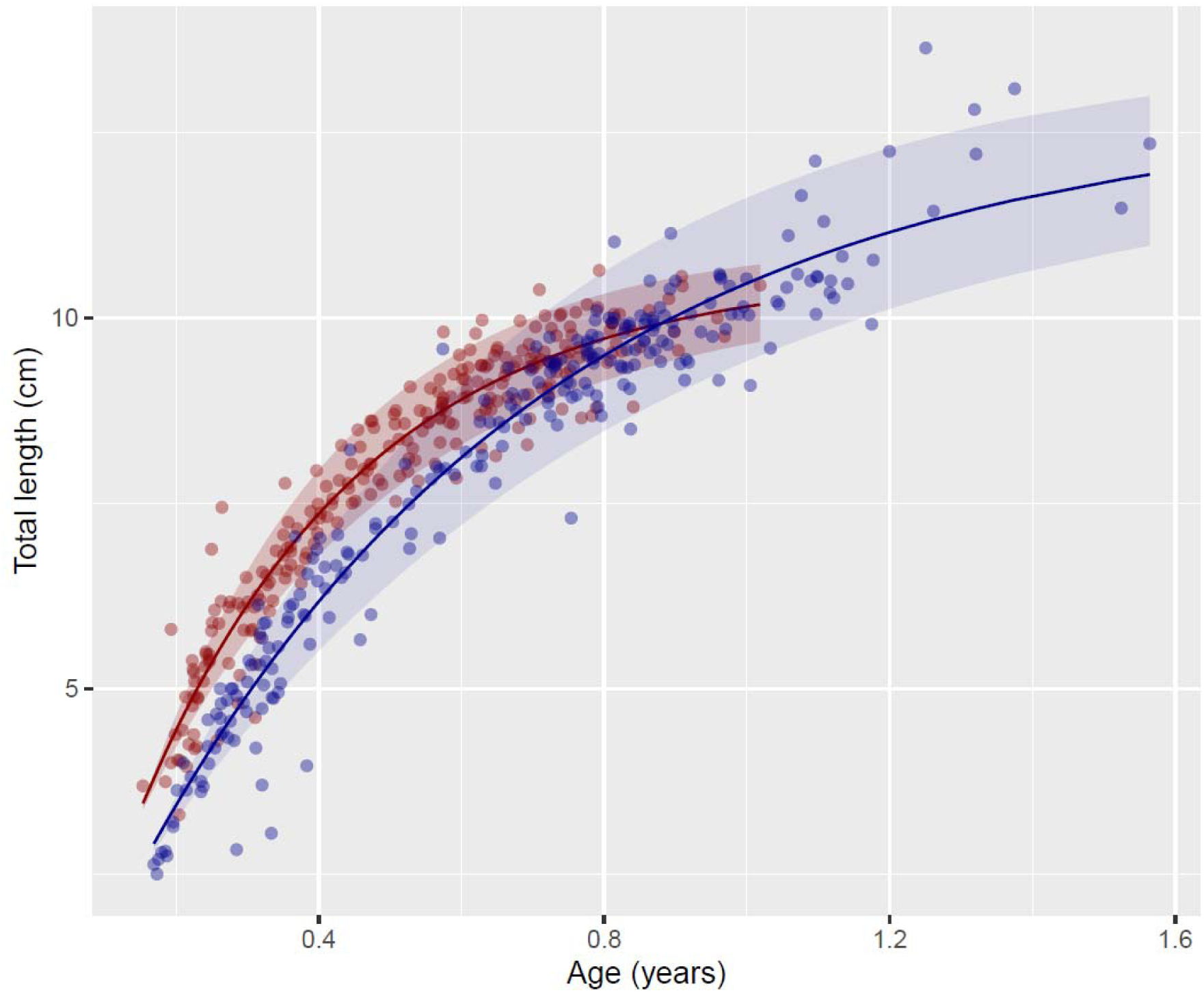
Von Bertalanffy growth functions (VBGF) for *Stolothrissa tanganicae* (in red) and *Limnothrissa miodon* (in blue), estimated from otolith-derived length-age relationships. Confidence intervals were obtained from growth curve estimations for the upper and lower limits of the 95 % confidence intervals of parameters L_∞_ and K.

## REFERENCES

Ahonen H. (2001). Age and growth of the pelagic clupeids in Lake Tanganyika estimated by otolith microstructure analysis. M.Sc. thesis, University of Turku, Finland.

Beamish R.J. & Mcfarlane G.A. (1983). The forgotten requirement for age validation in fisheries biology. Trans.Am. Fish. Soc. 112: 735–743.

Beverton R.J.H. & Holt S.J. (1957). On the dynamics of exploited fish populations. Gt Britain Fish Invest. Ser. 2, Vol. 19: 1-533.

Chapman D.W. & Van Well P. (1978). Growth and mortality of *Stolothrissa tanganicae*. Transactions of the American Fisheries Society 107: 26–35.

Chifamba C.P. (2019). The biology and impacts of *Oreochromis niloticus* and *Limnothrissa miodon* introduced in Lake Kariba. [Thesis fully internal (DIV), University of Groningen]. Rijksuniversiteit Groningen.

Cohen A.S., Bills R., Cocquyt C.Z. & Caljon A.G. (1993). Impact of sediment pollution on biodiversity in Lake Tanganyika. Conservation Biology 7: 667–677. 10.1046/j.1523-1739.1993.07030667.x

Coulter G.W. (ED.). (1991). Lake Tanganyika and its life. British Museum (Natural history) and Oxford University Press. London Oxford and New York. 354 pp.

De Keyzer E.L.R., De Corte Z., Van Steenberge M., Raeymaekers J.A.M., Calboli F.C.F., Kmentová N., Mulimbwa N., Virgilio M., Vangestel C., Mulungula P.M., Volckaert F.A.M. & Vanhove M.P.M. (2019). First genomic study on Lake Tanganyika sprat *Stolothrissa tanganicae*: a lack of population structure calls for integrated management of this important fisheries target species. Bmc Evolutionary Biology 19: 6. 10.1186/s12862-018-1325-8

De Keyzer E.L.R., Masilya Mulungula P., Alunga Lufungula G., Amisi Manala C., Andema Muniali A., Bashengezi Cibuhira P., Bashonga Bishobibiri A., Bashonga Rafiki A., Hyangya Lwikitcha B., Hugé J., Huyghe C.E.T., Itulamya Kitungano C., Janssens de Bisthoven L., Kakogozo Bombi J., Kamakune Sabiti S., Kiriza Katagata I., Kwibe Assani D., Lubunga Dunia P., Lumami KapepulaV., Lwacha F., Mazambi Lutete J., Shema Muhemura F., Milec L.J.M., Mulimbwa N’Sibula T., Mushagaulusa Mulega A., Muterezi Bukinga F., Muzumani Risasi D., Mwenyemali Banamwezi D., Kahindo N’djungu J., Nabintu Bugabanda N., Ntakobajira Karani J.-P., Raeymaekers J.A.M., Riziki Walumona J., Safari Rukahusa R., Vanhove M.P.M., Volckaert F.A.M., Wembo Ndeo O. & Van Steenberge M. (2020). Local perceptions on the state of fisheries and fisheries management at Uvira, Lake Tanganyika, Dr Congo. Journal of Great Lakes Research 46: 1740–1753. 10.1016/j.jglr.2020.09.003

Ehrenfels B., Junker J., Namutebi D., Callbeck C.M., Dinkel C., Kalangali A., Kimirei I.A., Mbonde A.S., Mosille J.B., Emmanuel A., Sweke E.A., Schubert C.J., Seehausen O., Wagner C.E. & Wehrli, B. (2023). Isotopic signatures induced by upwelling reveal regional fish stocks in Lake Tanganyika. Plos One 18(11): e0281828. 10.1371/journal.pone.0281828

Froese R. & Pauly D. (2017). FishBase [online]. [Cited 27 February 2024]. http://www.fishbase.org.

Gayanilo F.C. & Pauly D. 1997. Fao – ICLARM. Stock Assessment Tools ( FISAT). Fao Computerised information Series (Fisheries) 8, 262 p.

Junker J., Rick J.A., Mcintyre P.B., Kimirei I., Sweke E. A., Mosille J. B., Wehrli B., Dinkel C., Mwaiko S., Seehausen O. & Wagner C. E. (2020). Structural genomic variation leads to genetic differentiation in Lake Tanganyika’ s sardines. Molecular Ecology 29: 3277–3298. 10.1111/mec.15559

Kaningini M. (1995). Etude de la croissance, de la reproduction et de l’exploitation de *Limnothrissa miodon* au lac Kivu. Thèse de doctorat. Presse Universitaire de Namur (Belgique).

Kolding J., van Zwieten P., Marttin F., Funge-Smith S. & Poulain F. (2019). Freshwater small pelagic fish and fisheries in major African lakes and reservoirs in relation to food security and nutrition. *Fao Fisheries and Aquaculture* Technical Paper No 642. Rome, FAO. 124 pp.

Kimirei I. A. & Mgaya Y. D. (2007). Influence of environmental factors on seasonal changes in clupeid catches in the Kigoma area of Lake Tanganyika. African Journal of Aquatic Science 32: 291–298.

Kimura S. (1991B). Growth of *Stolothrissa tanganicae* estimated from daily otolith rings in southern Lake Tanganyika. In Kawanabe, H. and M. Nagoshi (eds), Ecological and limnological study on Lake Tanganyika and its adjacent regions, 7: 60–61.

Kimura S. (1991C). Growth of *Limnothrissa miodon* estimated from daily otolith rings in southern Lake Tanganyika. In Kawanabe, H. and M. Nagoshi (eds), Ecological and limnological study on Lake Tanganyika and its adjacent regions, 7: 62–64.

Kimura S. (1995). Growth of the clupeid fishes, *Stolothrissa tanganicae* and *Limnothrissa miodon*, in the Zambian waters of Lake Tanganyika. J. Fish Biol. 47: 569–575.

Kmentová N., Koblmüller S., Van Steenberge M., Raeymaekers J.A.M., Artois T., De Keyzer E.L.R., Milec L., Muterezi Bukinga F., Mulimbwa N., Mulungula P.M., Ntakimazi G., Volckaert F.A.M., Gelnar M. & Vanhove M.P.M. (2020). Weak population structure and recent demographic expansion of the monogenean parasite *Kapentagyrus* spp. infecting clupeid fishes of Lake Tanganyika, East Africa. International Journal for Parasitology 50: 471–486. 10.1016/j.ijpara.2020.02.002

Langenberg V.T. (2008). On the limnology of Lake Tanganyika. PhD. Thesis Wageningen University, The Netherlands. Isbn 978-90-8504-784-1.

Lta Secretariat (2012). Report on regional lake-wide fisheries frame survey on Lake Tanganyika 2011. LTA/Tech Doc/2012/01. Bujumbura, Burundi.

Lou D.C., Mapstone B.D., Russ G.R., Davies C.R. & Begg G.A. (2005). Using otolith weight-age relationships to predict age-based metrics of coral reef fish populations at different spatial scales. Fisheries Research 71: 279–294.

Mannini P., Aro E., Katonda I., Kassaka B., Mambona C., Milindi G., Paffen P. & Verburg P. (1996). Pelagic Fish Stocks of Lake Tanganyika: biology and exploitation. FAO/Finnida Research for the Management of the Fisheries of Lake Tanganyika. GCP/ RAF/271/Fin – TD/53 (En): 60 .

Mannini P. (1998). PhD. Thesis. Ecology of the Pelagic Fish Resources of Lake Tanganyika. University of Hull.

Mildenberger T. K., Taylor M. H. & Wolff M. (2017). TropFishR: an R package for fisheries analysis with length-frequency data. Methods in Ecology and Evolution 8: 1520– 1527). 10.1111/2041-210X.12791

Milec L.J.M., Vanhove M.P.M., Muterezi Bukinga F., De Keyzer E.L.R., Lumami Kapepula V., Masilya Mulungula P., Mulimbwa N’sibula T., Wagner C.E. & Raeymaekers J.A.M. (2022). Complete mitochondrial genomes and updated divergence time of the two freshwater clupeids endemic to Lake Tanganyika (Africa) suggest intralacustrine speciation. Bmc Ecology & Evolution 22: 127. 10.1186/s12862-022-02085-8

Meisfjord J., Midtøy F. & Folkvord A. (2006). Validation of daily increment deposition in otoliths of juvenile *Limnothrissa miodon* (Clupeidae). Journal of Fish Biology: 69: 1845–1848.

Moreau J., Munyandorera J. & Nyakageni B. (1991). Evaluation des paramètres demographiques chez *Stolothrissa tanganicae* et *Limnothrissa miodon* du lac Tanganyika. Verh . Internat. Verein. Limnol. 24: 2552 – 2558.

Mölsä H., Reynolds J.E., Coenen E. J. & Lindqvist O.V. (1999). Fisheries research towards resource management on Lake Tanganyika. Hydrobiologia 407: 1–24.

Mölsä H., Sarvala J., Badende S., Chitamwebwa D., Kanyaru R., Mulimbwa, N’sibula T., & Mwape L. (2002). Ecosystem monitoring in the development of sustainable fisheries in Lake Tanganyika. Aquatic Ecosystem Health and Management 5: 267–281.

Mulimbwa N’sibula T. & Shirakihara K. (1994). Growth, recruitment and reproduction of sardines (*Stolothrissa tanganicae* and *Limnothrissa miodon*) in Northwestern Lake Tanganyika. Tropics 4: 57–67.

Mulimbwa N’sibula T. (2006). Assessment of the commercial artisanal fishing impact on three endemic pelagic fish stocks, *Stolothrissa tanganicae*, Limnothrissa miodon and Lates stappersi in Bujumbura and Kigoma sub-basins of Lake Tanganyika. Verh. Internat. Verein. Limnol. 29: 1189–1193.

Mulimbwa N’sibula T., Sarvala J & Raeymaekers Jam (2014a). Reproductive activities of two zooplanktivorous clupeid fish in relation to the seasonal abundance of copepod prey in the northern end of Lake Tanganyika. Belgian Journal of Zoology 144: 77–92.

Mulimbwa N’sibula T., Raeymaekers J.A.M & Sarvala J. (2014b). Seasonal changes in the pelagic catch of two clupeid zooplanktivores in relation to the abundance of copepod zooplankton in the northern end of Lake Tanganyika. Aquatic Ecosystem Health & Management 17: 25–33.

Mulimbwa N’sibula T., Milec L.J.M., Raeymaekers J.A.M., Sarvala J., Plisnier P.-D., Marwa B. & Micha J.-C. (2022). Spatial and seasonal variation in reproductive indices of the clupeids *Limnothrissa miodon* and *Stolothrissa tanganicae* in the Congolese waters of northern Lake Tanganyika. Belgian Journal of Zoology 152: 13–31. 10.26496/bjz.2022.96.

Mulimbwa N’sibula T. (2020). Dynamique de *Limnothrissa miodon*, *Stolothrissa tanganicae* et de *Lates stappersi*, en vue de la gestion durable de la pêche au lac Tanganyika (sous bassin de Bujumbura). Rapport Mesurable, Rapportage et Vérifiable (MRV) CEBios 2020 Partenatiat Cooperation Belge au Développement (C.B.D)/ Institut Royal des Sciences Naturelles de Belgique, Bruxelles.

Mushagalusa C.D., Micha J.-C., Ntakimazi G. & Nshombo M. (2015). Brief evaluation of the current state of fish stocks landed by artisanal fishing units from the extreme northwest part of Lake Tanganyika. International Journal of Fisheries and Aquatic Studies 2: 41–48.

Okito M.G., Micha J.-C., Habarigura J. B., Ntakimazi G. & Nshombo M.V. (2017). Socio-economics of artisanal fisheries in Burundian waters of Lake Tanganyika at Mvugo and Muguruka. International Journal of Biological and Chemical Sciences, 11: 1991–8631.

O’reilly C.M., Alin S.R., Plisnier P.D., Cohen A.S. & Mckee B.A. (2003). Climate change decreases aquatic ecosystem productivity of Lake Tanganyika, Africa. Nature 424: 766–768.

Pauly D. & David N. (1981). Elefan I, a Basic program for the objective extraction of growth parameters from length-frequency data. Beriche der Deutschen Wissenschaftlichen Kommission für Meeresforschung 28: 205–211.

Plisnier P.-D. (1997). Climate, limnology, and fisheries changes of Lake Tanganyika. FAO/Finnida Research for the Management of the Fisheries of Lake Tanganyika. GCP/RAF/271/FIN-TD/72 (En):38p.

Plisnier P.-D., Mgana H., Kimirei I., Change A., Makasa L., Chimanga J., Zulu F., Cocquyt C., Horion S., Bergamino N., Naithani J., Deleersnijder E., André L., Descy J.P. & Cornet Y. (2009). Limnological variability and pelagic fish abundance (*Stolothrissa tanganicae* and *Lates stappersii*) in Lake Tanganyika. Hydrobiologia 625: 117– 134.

Plisnier P.-D., Kayanda R., MacIntyre S., Obiero K., Okello W., Vodacek A., Cocquyt C., Abegaz H., Achieng A., Akonkwa B., Albrecht C., Balagizi C., Barasa J., Abel Bashonga R., Bashonga Bishobibiri A., Bootsma H., Borges A.V., Chavula G., Dadi T., De Keyzer E.L.R., Doran P.J., Gabagambi N., Gatare R., Gemmell A., Getahun A., Haambiya L.H., Higgins S.N., Hyangya B.L., Irvine K., Isumbisho M., Jonasse C., Katongo C., Katsev S., Keyombe J., Kimirei I., Kisekelwa T., Kishe M., Otoung A. Koding S., Kolding J., Kraemer B.M., Limbu P., Lomodei E., Mahongo S.B., Malala J., Mbabazi S., Masilya P.M., McCandless M., Medard M., Migeni Ajode Z., Mrosso H.D., Mudakikwa E.R., Mulimbwa N., Mushagalusa D., Muvundja F.A., Nankabirwa A., Nahimana D., Ngatunga B.P., Ngochera M., Nicholson S., Nshombo M., Ntakimazi G., Nyamweya C., Ikwaput Nyeko J., Olago D., Olbamo T., O’Reilly C.M., Pasche N., Phiri H., Raasakka N., Salyani A., Sibomana C., Silsbe G.M., Smith S., Sterner R.W., Thiery W., Tuyisenge J., Van der Knaap M., Van Steenberge M., van Zwieten P.A.M., Verheyen E., Wakjira M., Walakira J., Ndeo Wembo O., Lawrence T. (2023). Need for harmonized long-term multi-lake monitoring of African Great Lakes. Journal of Great Lakes Research 49: 101988.

Rohatgi A. (2022). WebPlotDigitizer: Version 4.6. Pacifica, California, USA. https://automeris.io/WebPlotDigitizer

ROEST, 1978. Bibliography of fisheries and limnology for Lake Tanganyika. United Nations Food and Agriculture Organization, Cifa Occasional Paper, 6: 1–12 and supplements.

Sarvala. J. & Mulimbwa N’sibula T. (2023). GLOW10 2023. Dar es Salaam 15–17 February 2023.

Sarvala J., Langenberg V.T., Salonen K., Chitamwebwa D., Coulter G.W., Huttula T., Kanyaru R., Kotilainen P., Makasa L., Mulimbwa N. & Mölsä H. (2006A). Fish catches from Lake Tanganyika mainly reflect changes in fishery practices, not climate. Verh. Int. Ver. Theor. Angew. Limnol. 29: 1182–1188.

Sarvala J., Langenberg V.T., Salonen K., Chitamwebwa D., Coulter G.W., Huttula T., Kotilainen P., Mulimbwa N. & Mölsä H. (2006b). Changes in dissolved silica and transparency are not sufficient evidence for decreased primary productivity due to climate warming in Lake Tanganyika. Reply to comment by Verburg, Hecky and Kling. Verh. Int. Ver. Theor. Angew. Limnol. 29: 2339–2342.

Sarvala J., Salonen K., Järvinen M., Aro E., Huttula T., Kotilainen P., Kurki H., Langenberg V.T., Mannini P., Peltonen A., Plisnier P.-D., Vuorinen I., Mölsä H. & Lindqvist O. (1999). Trophic structure of Lake Tanganyika: carbon flows in the pelagic food web. Hydrobiologia 407:149–173.

Schwamborn R. (2018). How reliable are the Powell–Wetherall plot method and the maximum-length approach? Implications for length-based studies of growth and mortality. Reviews in Fish Biology and Fisheries 28: 587–605. 10.1007/s11160-018-9519-0

Schwamborn R., Mildenberger T.K. & Taylor M.H. (2019). Assessing sources of uncertainty in length-based estimates of body growth in populations of fishes and macroinvertebrates with bootstrapped ELEFAN. Ecological Modelling 393: 37–51. 10.1016/j.ecolmodel.2018.12.001

Sparre P. & Venema S.C. (1998). Introduction to tropical fish stock assessment, Part 1 - Manual. Fao Fisheries Technical Paper 306 (1): 407.

Taylor M.H. & Mildenberger T.K. (2017). Extending electronic length frequency analysis in R. Fisheries Management and Ecology 24: 230–238. 10.1111/fme.12232.

Van der Knaap M., Katonda I. & De Graaf G.J. (2014). Lake Tanganyika fisheries frame survey analysis: assessment of the options for management of the fisheries of Lake Tanganyika. Aquatic Ecosystem Health and Management 17: 4–13.

Van Steenberge M., Vanhove M.P.M., Muzumani Risasi D., Mulimbwa N’sibula T., Muterezi Bukinga F., Pariselle A., Gillardin C., Vreven E., Raeymaekers J.A.M., Huyse T., Volckaert F.A.M., Nshombo Muderhwa V. & Snoeks J. (2011). A recent inventory of the fishes of the north-western and central western coast of Lake Tanganyika (Democratic Republic Congo). Acta Ichthyologica et Piscatoria 41: 201–214. 10.3750/AIP2011.41.3.08

Verburg P., Hecky R.E. & Kling H. (2003). Ecological consequences of a century of warming in Lake Tanganyika. Science (80) 301: 505–507.

Von Bertalanffy L. (1938). A quantitative theory of organic growth. Hum. Biol. 10: 181 – 213

Wang K., Zhang C., Xu B., Xue Y. & Ren Y. (2020). Selecting optimal bin size to account for growth variability in Electronic LEngth Frequency ANalysis (ELEFAN). Fisheries Research 225: 105474. 10.1016/j.fishres.2019.105474

Wang K., Zhang C., Sun M., Xu B., Ji Y., Xue Y. & Ren Y. (2021). Fishing pressure and lifespan affect the estimation of growth parameters using ELEFAN. Fisheries Research, 238, 105903. 10.1016/j.fishres.2021.105903

